# Reconstructing co-dependent cellular crosstalk in lung adenocarcinoma using REMI

**DOI:** 10.1101/2021.05.02.440071

**Authors:** Alice Yu, Yuanyuan Li, Irene Li, Christine Yeh, Aaron E. Chiou, Michael G. Ozawa, Jonathan Taylor, Sylvia K. Plevritis

## Abstract

Cellular crosstalk in tissue microenvironments is fundamental to normal and pathological biological processes. Global assessment of cell-cell interactions (CCI) is not yet technically feasible, but computational efforts to reconstruct these interactions have been proposed. Current computational approaches that identify CCI often make the simplifying assumption that pairwise interactions are independent of one another, which can lead to reduced accuracy. We present REMI (REgularized Microenvironment Interactome), a graph-based algorithm that predicts ligand-receptor (LR) interactions by accounting for LR dependencies on high-dimensional, small sample size datasets. We apply REMI to reconstruct the human lung adenocarcinoma (LUAD) interactome from a bulk flow-sorted RNA-seq dataset, then leverage single-cell transcriptomics data to increase its resolution and identify LR prognostic signatures. We experimentally confirmed colocalization of CTGF:LRP6 as an interaction predicted to be associated with LUAD progression. Our work presents a novel way to reconstruct interactomes and a new approach to identify clinically-relevant cell-cell interactions.

## Introduction

Cell-cell interactions between and within the various cell types comprising the tissue microenvironment play a fundamental role in regulating local and systemic biological and physiological functions under normal and pathological conditions. These interactions facilitate cooperation or competition between cell types and are typically mediated between ligands and receptors. Ligands are often manifested as soluble or extracellular proteins that are expressed by the “sending” cells and bind onto a cognate receptor on the “receiving” cells (*1*). In tumor microenvironments (TME), cellular crosstalk between tumor, stroma, and immune cells orchestrates the establishment of pre-invasive and invasive niches that enable cancer progression properties, such as tumor growth, immune evasion, and metastasis. Large-scale cellular interactions in these milieus are difficult to measure using current experimental techniques, but several computational approaches have been proposed to predict these interactions using -omics data.

A majority of the current computational approaches that utilize high-throughput transcriptomics data to infer cell-cell interactions either calculate interaction scores based on gene expression permutation tests or implement graph-based approaches (*2*). Expression permutation-based approaches, such as CellphoneDB and NATMI, threshold ligand and receptor genes based on their expression values to identify cell type-specific ligand and receptors with the assumption that this predicts higher ligand-receptor (LR) protein abundance (*3, 4*). Methods, such as CCCExplorer, calculate correlation metrics between the expression levels of the ligand, receptor, and downstream signaling pathway genes for each LR pair (*5*). However, a correlation between the expression of ligand and receptor genes may be capturing an indirect association caused by another LR interaction involving the ligand or receptor of interest. NicheNET, instead, predicts what downstream target a ligand is most likely associated with using metrics calculated on prior knowledge graphs rather than solely relying on the experimental data (*6*), but does not specify implicated receptors.

While current crosstalk inference approaches provide a valuable baseline for synthesizing hypotheses of LR interactions, they do not capture the direct vs indirect dependencies of LR pairs within the TME. Current approaches make the assumption that pairwise interactions are independent, but a pairwise interaction can be influenced by other interactions via autocrine and paracrine loops (*7, 8*). Calculating dependent versus independent LR pairs is often an ill-defined problem for high-dimensional datasets. To address this challenge, we present the algorithm called REMI (REgularized Microenvironment Interactome), to identify communities of dependent LR pairs on high-dimensional datasets of relatively small sample size using graph-based approaches. We demonstrate the performance of REMI by simulating datasets with varying sample sizes to show how REMI outperforms existing approaches in datasets with small sample size.

To compare REMI to existing interactomes, we focused on reconstructing the lung adenocarcinoma (LUAD) interactome. Various renditions of the LUAD microenvironment have been assembled using different approaches that have led to novel insights. Kumar et al. built an interactome for LUAD using mouse single-cell RNA-seq (scRNA-seq) data, where they used a scoring mechanism that captured highly expressed LR genes (*9*). Gentles et al. created the Lung Tumor Microenvironment Interactome (LTMI) from bulk flow-sorted RNA-Seq data by thresholding gene expression levels in the dataset and computing pairwise correlations between ligand and receptor genes (*10*). We applied REMI to the LTMI dataset to recreate a more specific rendition of LUAD interactome (REMI-LUAD). We then projected an independent LUAD scRNA-seq dataset onto REMI-LUAD to increase the resolution of the interactome and referred to this interactome as the single-cell rendition of REMI LUAD (scREMI-LUAD). Using scREMI-LUAD, we identified paracrine signaling interactions between subpopulations of a given cell type that were previously annotated as autocrine signaling interactions. To generate a signature of LUAD progression from scREMI-LUAD, we assigned a prognostic score to each cell subpopulation and identified prognostically-associated crosstalk signatures that may lead to clinically-relevant biomarkers and therapeutic targets. In summary, REMI offers a new approach to reconstruct cell-cell crosstalk underlying the tumor microenvironment by accounting for conditional dependencies using graph-based approaches. REMI is implemented in R and is freely available on GitHub (https://github.com/ayu1/REMI).

## Results

### Impact of correlation versus partial correlation analysis on ligand-receptor pair inference

Many of the current computational approaches use correlation to identify potential occurring LR interactions. Here, we show that correlation analyses result in false positive predictions whereas measuring the partial correlation of LR pairs increases specificity. Partial correlation, similar to regression, removes potential confounding factors from correlation values that are caused by another ligand or receptor indirectly affecting the expression levels of the LR pair of interest. We started by obtaining 30 non-small cell lung carcinoma primary specimens and focused on analyzing 17 lung adenocarcinoma (LUAD) primary specimens from a publicly-available bulk flow-sorted RNA-seq dataset, which contains gene expression for malignant, fibroblast, endothelial, and pan-immune cells (GSE111907) (*10*). Hierarchically clustering on only the genes that expressed ligands and receptors grouped the samples by cell type (Fig 1a). We then constructed a LR correlation network using the data (*G_LTMI_*), where the nodes represent ligand and receptor genes and an edge represents a LR pair using the threshold criteria set by Gentles et al. The edge weight is set as the Pearson correlation between the gene expression of the ligand and receptor.

**Figure 1.**
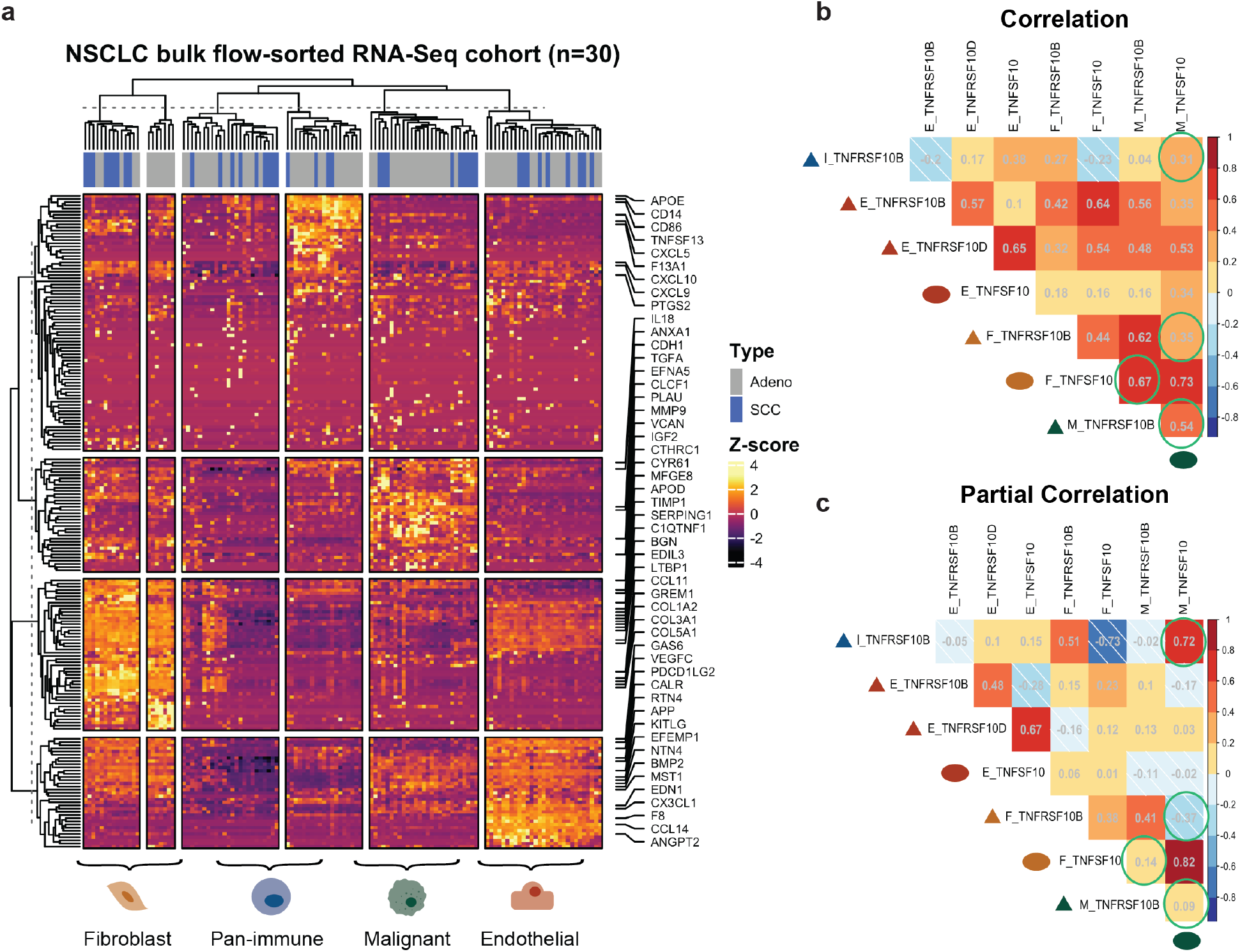
Network representation of Lung TMI shows dense components of cognate ligand and receptor pairs. **(a)** All ligand and receptor genes in the LUAD flow-sorted RNA-Seq dataset hierarchically clustered. Genes labeled are the most highly expressed genes in the top five percentile. **(b)** Correlation and partial correlation values of the nodes that are found within the circled component in the LTMI network. Calculated correlation matrix values are on the left and partial correlation matrix values are on the right. The text labels of the correlation matrices are the gene’s cell-type added in front of the gene name (celltype_name). Additionally, a colored (cell type) shape (gene function) is placed in front of the text label. Colors correspond to legend in (b). Bottom dark green circle corresponds to M_TNFSF10 labeled at the top of the column. Green circles highlight examples of correlation values that are changed after accounting for confounding variables. White stripes indicate a negative partial correlation value.

Since many receptors have multiple ligand pairings and vice versa, the correlation network is composed of multiple dense and disjoint network components. To infer dependent LR pairs within the network, we calculated the partial correlation of genes on a small disjoint subgraph, which contains one ligand node (*TNFSF10)* and two receptor nodes (*TNFRSF10B*, *TNFRSF10D)* across multiple cell types (Supp Fig 1a) (Fig 1b). Simultaneous TNFSF10:TNFRSF10B and TNFSF10:TNFRSF10D interactions could occur, but one interaction plays an agonist role while the other plays an antagonistic role (*11*). When TNFSF10 binds onto TNFRSF10B, it induces cell apoptosis via the TRAIL pathway, whereas TNFSF10 inhibits the TRAIL pathway when it binds onto TNFRSF10D (*12*). If we infer LR interactions based on correlation, the ligand TNFSF10 secreted from the malignant cell will interact with the receptor TNFRSF10B on malignant and fibroblast cells (circled in green), indicating likely apoptosis of malignant cells and cancer-associated fibroblasts (Fig 1c). This contradicts studies that show pro-tumoral effects of fibroblasts and how they can express the decoy receptor to avoid apoptosis (*13*). Instead, by calculating the partial correlation of the LR pairs within the circled component, we find more biologically-reasonable results. Based on the partial correlation values, TNFSF10 expressed in the malignant cells is more likely to interact with TNFRSF10B expressed in the immune cells, as described in literature (*12, 14*). This simple analysis demonstrates how partial correlation removes confounding effects in correlation analysis and can thereby reduce false positive edges when building the interactome (right of Fig 1b). However, calculating the partial correlation is infeasible on larger subgraphs due to high-dimensionality of the dataset. REMI extends this concept onto larger components of the network.

### REMI algorithm

To remove confounding effects in LR networks, we introduce our novel algorithm, REMI, which identifies highly specific communities of conditionally-dependent cell-cell interactions on high-dimensional transcriptomic datasets with small sample sizes. REMI is comprised of four steps: (i) build a weighted undirected LR correlation network leveraging known LR pairings, (ii) detect communities of LR groups, (iii) identify conditionally-dependent LR pairs on communities, and (iv) reconstruct the resulting interactome (Fig 2a) (Methods). An additional step in REMI allows for the user to measure the significance of a LR pair prediction with respect to the LR pair’s REMI community.

**Figure 2.**
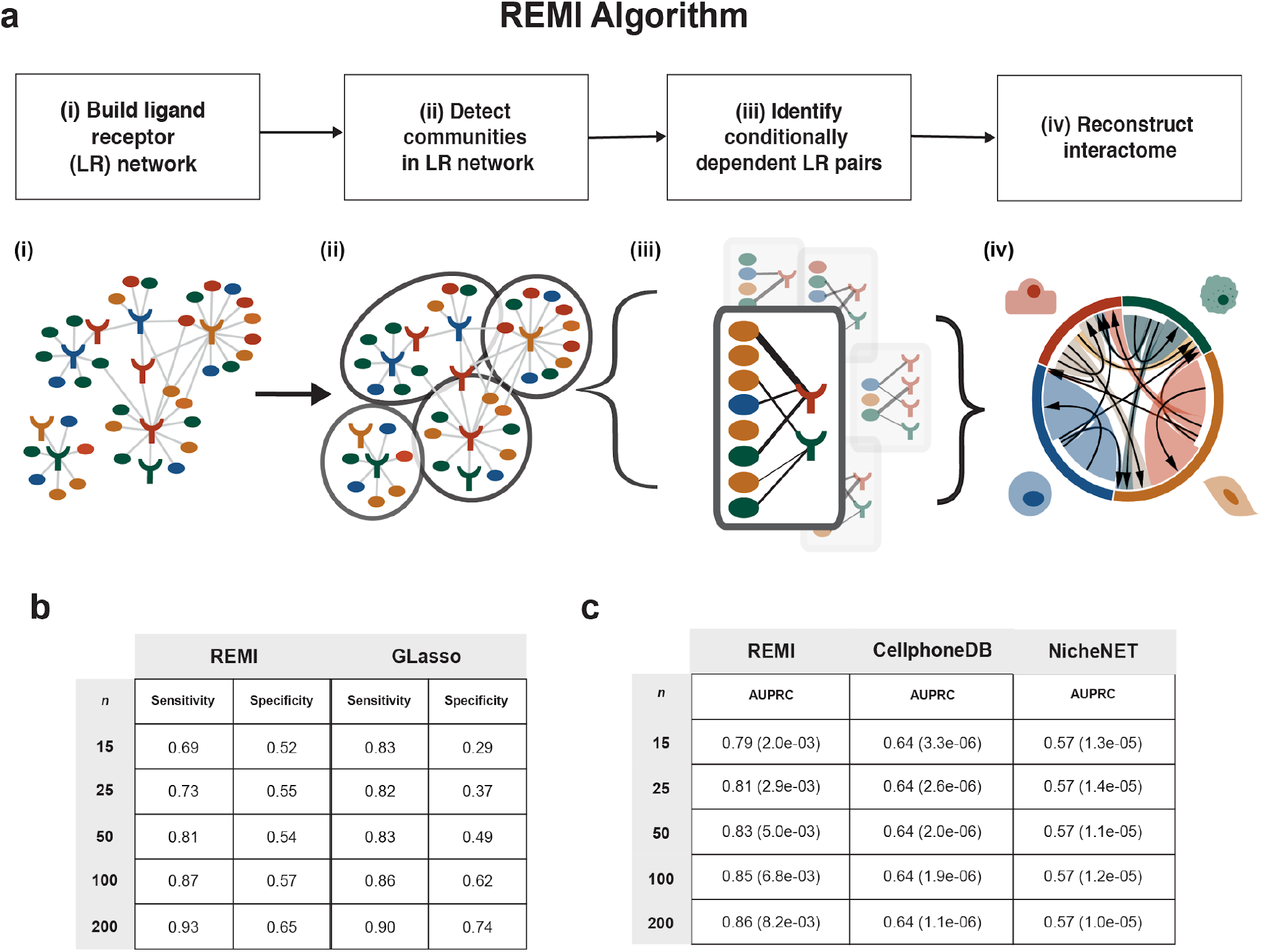
General REMI workflow and evaluation. **(a) (i)** A network of all possible LR pairs in TME is built. Nodes represent either ligand or receptor genes. Edges represent known LR pairs. **(ii)** Communities are identified within the LR network based on density **(iii)** Each community is converted into a graphical Gaussian model (GGM) and conditionally independent LR pairs are removed **(iv)** Conditionally dependent LR pairs from each community are reassembled to construct the global interactome. **(b)** Measuring the performance of REMI versus GLasso using a simulated gold standard interactome calculated from the LUAD TCGA RNA-seq dataset. Within the TCGA dataset, various cohort sizes were sampled to test REMI’s performance. Each cohort size was randomly sampled 50 times. **(c)** Test performance of REMI compared to other computational crosstalk prediction algorithms using our TCGA gold-standard interactome. Each cohort size was randomly sampled 50 times. Value in parentheses is the standard deviation of AUPRC. **(b)** Force-directed network representation of LTMI (V = 868, E = 2652). Nodes represent ligand (circle) and receptor (triangle) genes and an edge represents a predicted ligand receptor pair. Nodes are colored by cell type.

In its first step, REMI generates a weighted bipartite LR network, denoted by *G*, where the nodes represent either a ligand or receptor gene expressed in a specified cell type within the dataset. Edges of *G* are drawn between literature-supported LR pairings curated from the FANTOM5 database (*15*). Edge weights of *G* are computed as the Pearson correlation between the gene expression of the ligand and receptor nodes (Fig 2a, i). Ideally, the conditional-dependency of each LR pair would be computed with respect to all other nodes in the network. However, current high-dimensional transcriptomic datasets contain a large number of genes compared to the number of samples, which makes the conditional-dependency hard to estimate with high accuracy.

To reduce the size of the LR network, REMI hierarchically divides *G* into groups of densely connected nodes, called communities, using the Louvain community detection method (*16*). Next, REMI identifies conditionally-dependent LR pairs in each community independently. An inverse-covariance matrix is created for each community using the data measuring the covariance between all genes within the community. The resulting matrix represents the partial correlation of a LR pair with respect to other pairs in the community (Fig 2a, ii). We set edges between ligands and edges between receptors as zero. Then the community network is regularized using graphical lasso (GLasso) to identify conditionally-dependent LR edges (Fig 2a, iii) (*17*). These edges represent LR pairs that have a strong association, while accounting for correlated LR pairs within their community that may have a confounding effect. The resulting edge weights represent the regularized partial correlation metrics of the LR pair with respect to its community. For further downstream analyses, we analyzed a binarized version of the network, denoted as 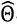, reconstructed by aggregating the communities after filtering for edges with a positive edge weight.

The community detection algorithm component of REMI defines what LR pairs may be confounding one another. We ran several simulations to test the sensitivity and specificity of varying community sizes. REMI’s best performance occurs when the number of nodes in each community is equal to the sample size. Simulation details will be elaborated in the next section (Supp Figure 1C, Supplementary Materials). Therefore, REMI hierarchically breaks down large communities until the size of the communities is approximately equivalent to sample size. Communities identified by Louvain are referred to as “within-communities” (WC). This step prioritizes assigning a node to a unique community, but leaves edges that represent potential LR pairs unassigned. REMI creates additional communities to account for the newly unassigned edges by aggregating unassigned edges that exist between two communities to create “between-communities” (BC) (Supp Fig 1a). The conditionally-dependent WC and BC edges are aggregated to reconstruct the interactome (Fig 2a, iv).

To account for the independent community calculations, we developed a way to measure the significance of each computed edge with respect to its community. The p-value is calculated by randomly permuting the edge-weight of the LR pair with randomly sampled correlation values. Since multiple correlation values can predict a binary edge in REMI, we build a decision tree to sort out which correlation values predict the edge of interest. We then sample from our null Wishart distribution, which represents a prior covariance matrix distribution, and predict edges using the decision tree. Edges predicted using values sampled from the null represents our REMI null distribution. We then calculate a one-sided p-value by measuring the number of null edges that had a greater correlation value than the original edge weight. The one-sided p-value is then converted to a two-sided p-value (Supp Fig 1b). Although the statistical test is computationally intensive, it can be used for determining the presence or absence of the edges when aggregating the communities to reconstruct the global network.

### Testing robustness of REMI parameters via simulations

Experimentally measured global interactome networks do not yet exist. To assess REMI’s performance in recovering LR predictions for a relatively small sample size dataset, we generated a bulk-level interactome by leveraging the large publicly available Cancer Genome Atlas (TCGA) LUAD bulk RNA-sequencing dataset (*n* = 1013). We made the assumption that the 1013 patients in this dataset represent an entire population. We then constructed *G* from the TCGA dataset (*V* = 893, *E* = 1331) and regularized *G* using the Graphical Gaussian model approach, GLasso, to estimate the conditionally-dependent network, denoted as 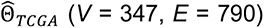, which we assume as our gold-standard (Supp Dataset). This represents a network of conditionally-dependent LR pairs, where the confounding effect of all possible interactions within the microenvironment is accounted for. From the TCGA LUAD cohort, we created cohorts of different sample sizes (*n* = 15, 25, 50, 100, 200), randomly sampling the TCGA samples fifty times each respectively. We ran REMI and GLasso on each sampled dataset and compared their performance in terms of sensitivity and specificity (Fig 2b – c).

For large sample sizes datasets (n=200), REMI and GLasso have relativity high sensitivity and specificity, as expected (Fig 2b). As the sample size decreased from 200 to 15 patients, GLasso retained high sensitivity values compared to REMI. GLasso’s sensitivity dropped from 0.90 to 0.83, whereas REMI’s sensitivity dropped from 0.93 to 0.69. However, REMI retained higher specificity than GLasso as the sample size decreased: GLasso’s specificity dropped from 0.74 to 0.29 (61%), whereas REMI’s specificity dropped from 0.65 to 0.52 (20%). In summary, REMI has higher specificity compared to GLasso, which we regard as a preferred property when selecting candidate-relevant interactions for experimental validation.

To further dissect the performance of REMI, we tested the performance of within-community (WC) and between-community (BC) predictions as a function of sample size on the TCGA simulation analysis. WCs have a higher density compared to BCs and yielded relatively higher sensitivity and lower specificity as sample size decreased from *n* = 200 to *n* = 15 (Supp Fig 1c). In WCs, sensitivity dropped from 0.93 to 0.75 (18% decrease) and specificity dropped from 0.63 to 0.45 (18% decrease). On the other hand, BCs performed with relatively lower sensitivity and higher specificity as sample size decreased from *n* = 200 to *n* = 15. In BCs, sensitivity dropped from 0.88 to 0.45 (43%) and specificity stayed relatively the same from 0.74 to 0.76 (2% increase)(Supp Fig 1c). Next, we considered the effect of the community size on REMI’s performance. When the number of nodes within a community was increased 2*n* to 50*n* times, the density of the communities increased and specificity decreased from 0.52 to 0.33 (19%) whereas sensitivity increased from 0.66 to 0.78 (12%) (Supp Fig 1d). Community sizes closer to the sample size of the dataset have increased specificity as opposed to larger community sizes. This further confirms that REMI optimizes for specificity over sensitivity.

### Comparison of REMI to other approaches

We compared REMI to current computational approaches designed for bulk RNA-seq and scRNA-seq that infer cell-cell interactions. We selected methods that represent the expression permutation-based approaches (Cellphone DB v2.0 and NATMI) and network-based approaches (CCCExplorer and NicheNET). Using the TCGA dataset, we ran the methods on the same sampled cohorts and used (Fig 2d). NicheNET performed on average an AUC of 0.65 across all cohort sizes and CellphoneDB v2.0 performed on average an AUC of 0.55 across all sample sizes. The difference in performance metrics can be explained by the difference in the assumptions each algorithm makes. CellphoneDB v2.0 predicts LR pairs based on the mean, however this can be a weak predictor in small sample sizes due to high variability across patients. NicheNET relies both on literature-derived models and user-defined inputs to specify downstream-activated genes. Since similar genes were listed as downstream genes across the different sampled datasets, NicheNET generated similar results. For NATMI, the method’s scoring system is dependent on the comparison of one cell-type’s expression with respect to other cell-types. Since we can only capture autocrine signaling interactions in this bulk dataset simulation, NATMI did not return comparable results. For CCCExplorer, the method is reliant on a control and experimental group, which differs from the assumptions made by the simulation. Other approaches were designed solely for single-cell RNA-seq and we compared those approaches to our results in later analyses. Along the spectrum of the bias-variance trade-off, our simulation analysis shows that current algorithms skew towards the bias-end, whereas REMI skews towards the variance-end, better capturing signals within the data to make predictions.

### Reconstructing LUAD interactome using REMI

We applied REMI to the bulk flow-sorted LUAD RNA-Seq cohort to assemble the REMI-LUAD interactome. REMI has 16% fewer nodes compared *G_LTMI_* which was constructed using the dataset in the LTMI resource (*V* = 868 in LTMI, *V* = 727 in REMI) and 45% fewer edges (*E* = 2652 in LTMI, *E* = 1435 in REMI), highlighting the specificity of REMI’s approach (Fig 3a)(Supplementary Dataset). REMI-LUAD removed LR edges that were densely connected (average node degree of edges removed = 12) and also identified a few new LR pairs (Supp Fig 1c). In REMI-LUAD, known lung cancer-specific LR pairs were identified, including *GREM1:KDR* which was experimentally validated between fibroblast and malignant cells (*10*)(Fig 3b). We observed fewer immune interactions within REMI-LUAD than LTMI because we suspect greater heterogeneity among the pan-immune cells relative to the other cell types in the bulk-sorted samples, as demonstrated in many published single cell RNA-seq studies (*18*).

**Figure 3.**
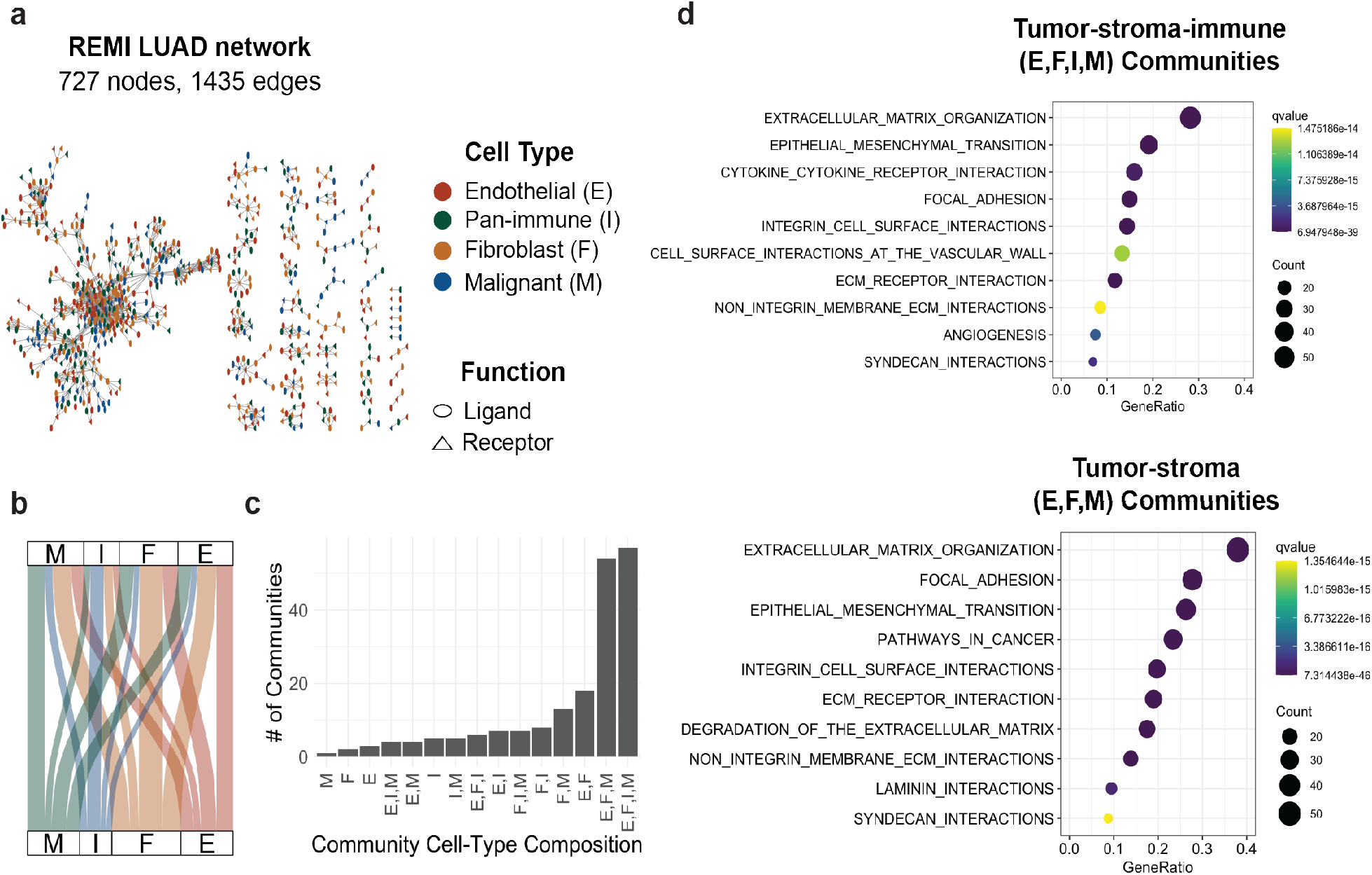
Reconstructed LUAD tumor microenvironment interactome reveals tumor-stroma specific communities. **(a)** Force-directed layout of REMI-LUAD interactome network. Shape of the nodes represent gene function. Color of the node represents cell type as denoted by the legend. **(b)** Alluvial plot showing ratio of paracrine and autocrine signaling interactions occurring between cell types. Thickness of lines represent the number of LR pairs. Full table is available in the supplements. **(c)** Distribution of number of communities with certain cell-type compositions **(d)** Top ten most significant enrichments for genes within the tumor-immune-stroma communities and tumor-stroma communities. GeneRatio is the ratio of the number of community-specific genes that overlapped with the geneset to the number of community-specific genes that overlapped with all the genesets in the collection. Count represents the number of genes.

To understand which cell types are interacting in REMI-LUAD, we analyzed the cell-type composition within each community. Majority of the communities were comprised of either all cell types (E, F, I, M) (36%) or tumor-stroma cells (E, F, M) (25%) (Fig 3c). To gauge the phenotypic properties enriched in these immune-stroma and stroma communities, we performed geneset enrichment analysis (GSEA) on the LR genes in the community groups. Communities with tumor-immune-stroma interactions were enriched for cytokines, angiogenesis, extracellular matrix (ECM) organization, and leukocyte extravasation. Angiogenesis and ECM remodeling are crucial for building a niche architecture for tumor growth. Communities with tumor-stroma specific interactions were uniquely enriched for degradation of the ECM (Fig 3d). ECM stiffness increases immunosuppression and stimulates epithelial transformation in the tumor, whereas ECM degradation aids in creating paths for cancer cell migration (*19*). Integrins found within REMI-LUAD, in particular, can detect ECM mechanical stiffness and assist with cell migration through degraded areas (*20*). Our results suggest that REMI-LUAD captures cell-cell interactions associated with processes ranging from tumor growth to invasion.

### Increasing cellular resolution of REMI-LUAD using a scRNA-seq LUAD dataset

In order to gain higher cellular resolution of the REMI-LUAD, we analyzed REMI-LUAD using a publicly-available independent scRNA-seq LUAD dataset from Lambrecht et al. (two patients, 22,681 cells) (E-MTAB-6149) (*21*). Using the annotated cell-type labels from the manuscript, we re-clustered them individually using Louvain clustering (Fig 4a). We labeled the immune clusters with broad immune cell subtypes as denoted from the manuscript (Myeloid, T cells, and B cells). For the remaining cell types, we identified five malignant (denoted as M(1), M(2), M(3), M(4), M(5)), three endothelial (denoted as E(1), E(2), E(3)) sub-populations, and four fibroblast subpopulations (denoted as F(1), F(2), F(3), F(4)). The four fibroblast subpopulations matched the ones identified by Lambrecht et al. and were further labeled as myofibroblasts (MyoFibs), inflammatory fibroblasts (iFibs), pericytes, and fibroblasts (Fibs) respectively based on their cell subpopulation-specific marker genes (Fig 4b – d) (Supp Fig 3a) (*22*).

**Figure 4.**
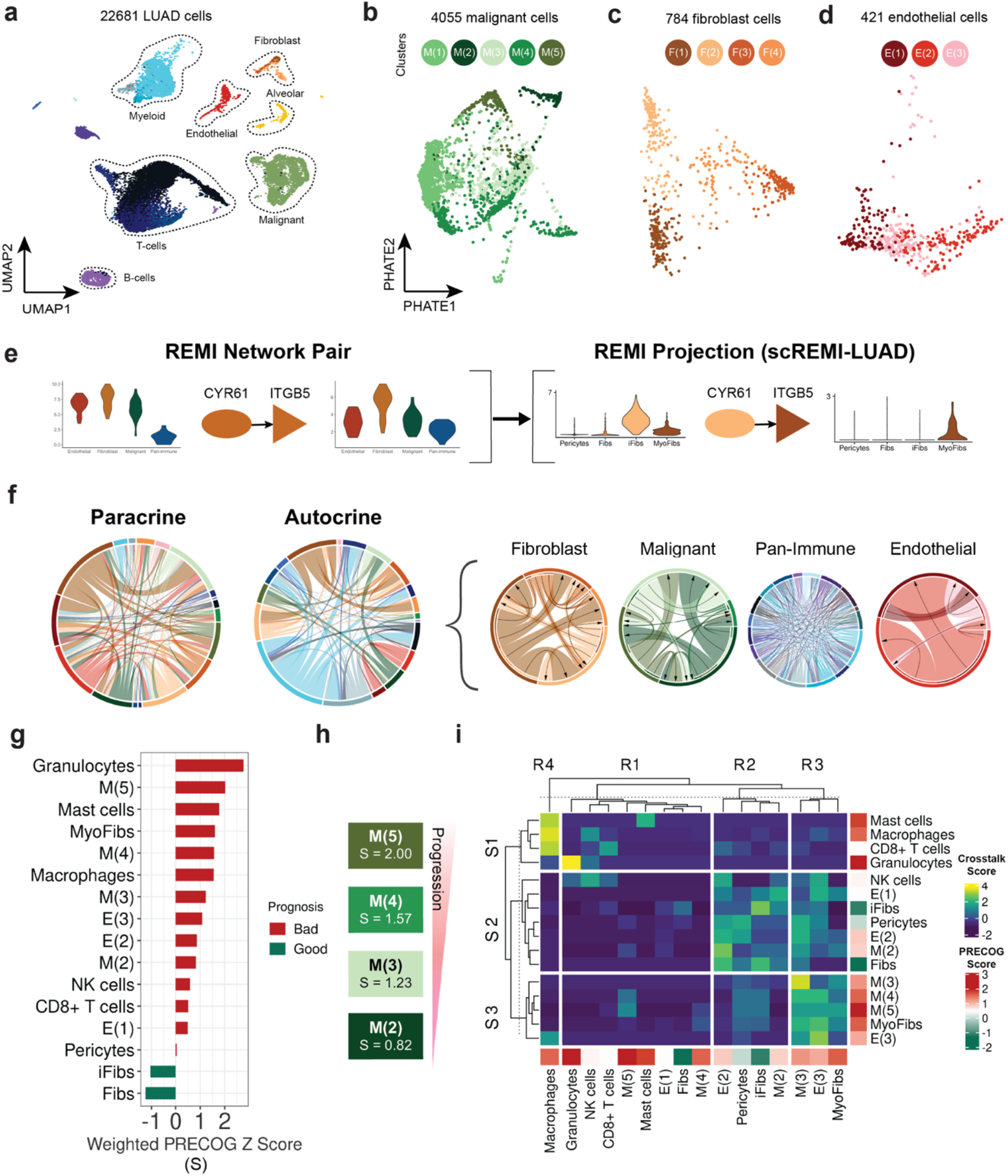
Projecting single cell RNA-sequencing information onto LUAD interactome reveals prognostic progression in malignant subpopulations. **(a)** Uniform manifold approximation and projection (UMAP) of Lambrecht et al. lung carcinoma single-cell RNA-Seq dataset (n=22681 cells, 2 patients) **(b)** Potential of heat diffusion for affinity-based transition embedding (PHATE) representation of malignant cell population and its five subpopulations (*n*=4055 cells) **(c)** PHATE representation of fibroblast population. Four subpopulations for fibroblast cells categorized as myofibroblasts (Fib(1)), inflammatory fibroblasts (Fib(2)), pericytes (Fib(3)), and fibroblasts (Fib(4)) (*n* = 784 cells) **(d)** PHATE representation of endothelial population and its three subpopulations (*n* = 421 cells) **(e)** Deconvolution pipeline of REMI interactome using single cell RNA-seq data. The example shows one LR pair where its cell-type node label is re-labeled by the single-cell subpopulation labels. (**f)** Proportion of interactions involving each cell type in scREMI-LUAD. Two chords on the left show interactions that were labeled as paracrine and autocrine in REMI-LUAD. Four chords on the right show the interactions that were previously labeled as autocrine are now labeled as paracrine signaling interactions between subpopulations. **(g)** Weighted prognosis scores for each subpopulation **(h)** Progression of malignant subpopulations ranked from poor to good prognosis. **(i)** Clustered cell-cell interaction crosstalk scores between cell sub-populations. Crosstalk score represents the variability of number of LR interactions between two cell types with respect to the sending cell type. Cell types are labeled by weighted prognosis score.

To adapt REMI for scRNA-seq data, we averaged each gene’s expression across each cell-type per patient. The Lambrecht et al. dataset consists of only two LUAD patients, which does not provide enough power for REMI’s correlation-based calculations. For this reason, we simply projected the scRNA-seq cell types onto REMI-LUAD and relabeled the nodes’ cell types based on the expression levels in the single-cell cohort. This network is referred to as single-cell REMI-LUAD (scREMI-LUAD). For each subpopulation, we filtered for differentially expressed (DE) ligands and receptors that had an averaged expression level greater than 0.4 (Supp Fig 3 a – c). Reassuringly, we found that 65% of the interactions in REMI-LUAD were present in scREMI-LUAD. Because many LR genes appear in multiple sub-populations, number of interactions in scREMI-LUAD is greater than REMI-LUAD, providing potential insight into the extent of heterogeneity of the interactome. For example, REMI-LUAD inferred that fibroblasts secrete *CYR61*, which binds onto *ITGB5* on fibroblast cells via an autocrine loop. In scREMI-LUAD, *CYR61* is expressed in iFibs and *ITGB5* in myofibroblasts. Hence the REMI-LUAD *CYR61:ITGB5* autocrine fibroblast interaction is now defined as a paracrine signaling between different subpopulation of fibroblasts in scREMI-LUAD (Fig 4e). Notably, many autocrine signaling interactions were converted into paracrine interactions between subpopulations of a given cell type (Fig 4f). The increased amount of paracrine intercellular crosstalk highlights the extent of the complex crosstalk between TME subpopulations.

### Inferring cellular progression in scREMI-LUAD by leveraging the prognostic significance scores in each subpopulation

To determine whether the cooperation between subpopulations is involved with disease progression, we calculated a score representative of each subpopulation’s ligand prognosis signature. We downloaded LUAD-specific prognostic gene scores from the PRECOG (PREdiction of Clinical Outcomes from Genomic profiles) database, which were calculated from bulk survival meta-analysis (*23*). A positive PRECOG z-score is associated with poor prognosis and a negative PRECOG z-score is associated with good prognosis. To assign the scores to each subpopulation in scREMI-LUAD, a weighted prognosis score was calculated by multiplying each differentially expressed ligand’s average cell-type expression by its PRECOG score in each subpopulation. We then aggregated the weighted ligand scores for each subpopulation and normalized it by the sum of all the ligand’s expression levels (Fig 4g). Granulocytes, MyoFibs, M(3), M(4), and M(5) expressed a poor-prognostic ligand signature, whereas iFibs expressed a good-prognostic ligand signature. While the malignant subpopulations all have poor-prognostic scores, the scores can be ordered, suggesting that different malignant cell subpopulations may be associated with different phases of tumor progression (Fig 4h). From this perspective, we filtered scREMI-LUAD by keeping the ligand from the subpopulation that had the closest weighted PRECOG score to its receptor in order to focus on interactions potentially associated with disease progression. We then calculated the crosstalk score by calculating the number of interactions occurring between each subpopulation. Then, we standardize the count with respect to the number of interactions involving the sending cell. This measures the variability of the number of interacting ligands for each sending cell subpopulation.

Crosstalk scores were clustered using k-means consensus clustering and three groups of sending (S1, S2 and S3) and four groups of receiving (R1, R2, R3 and R4) subpopulations (Fig 4i) were identified. Interestingly, S2 and R2 are enriched for subpopulations associated with relatively good prognosis, such as iFibs, pericytes, E(1), E(2), NK cells, and M(2). S3 and R3 are enriched for subpopulations associated with poor prognosis, including the remaining malignant subpopulations, E(3), and MyoFibs. We highlight three clusters of scores: (i) good-prognostic interactions between S2 and R2 (ii) mixed-prognostic interactions between S2 and R3, and (iii) poor-prognostic interactions between S3 and R3. To understand what types of interactions are occurring within each group, we performed GSEA on the good, mixed, and poor prognostic interactions (Supp Fig 4a–c). The good prognostic interactions are enriched for coagulation, elastic fiber formation, and more. One of the interactions occurs between SLIT2:ROBO1 as an autocrine interaction amongst inflammatory fibroblasts. Interestingly, Slit/Robo signaling has been shown to inhibit lung cancer migration in murine and *in-vitro* models (*24*, *25*) and we suspect it may play a similar role in primary lung cancer. The mixed-prognostic interactions are enriched for angiogenesis, ECM degradation, platelet adhesion to collagen, and more. One interaction occurring between M(2) and M(3), MMP7:CD151, has been confirmed to interact at the edge of LUAD nests. This suggests that mixed-prognostic interactions may occur on the invasive front (*26*). The poor-prognostic interactions were enriched for ECM degradation, angiogenesis, Notch signaling activation, and more. Interestingly, these interactions involve the ligands Tenascin-C and Versican, which are associated with metastatic phenotypes, such as cancer stem cell maintenance and mesenchymal-to-epithelial transition (*27, 28*). Based on the enrichments of interactions within each group, we reason that good prognostic interactions may be associated with noninvasive TMEs, poor prognostic interactions may be associated with invasive TMEs, and mixed-prognostic interactions may capture the transition from a noninvasive to invasive TMEs.

### *In-situ* experimental validation of CTGF and LRP6 co-localization among malignant subpopulations in LUAD

We focused on the mixed-prognostic interactions because these interactions may be involved with priming the tumor microenvironment for invasion. Proportionally, many mixed-prognostic interactions occur between the malignant subpopulations M(2) and M(3), suggesting potential subclonal cooperation between these subpopulations (Fig 5a). Among the M(2) to M(3) interactions, we highlight the LR interaction, CTGF:LRP6, due to the increasing interest in anti-CTGF therapy (Fig 5b) (*29, 30*). CTGF is inferred to be secreted not only from M(2), but also iFibs from within the same scREMI-LUAD community (p-val < 0.1)(Fig 5c and Supp Fig 5a-b). LRP6 is a co-receptor for the WNT signaling pathway and regulates tissue homeostasis (*31*). CTGF is a complex gene involved in many different processes, such as wound healing, angiogenesis, cell adhesion, migration, fibrosis, and ECM deposition (*32*). It activates both TGF-*β* and WNT/*β*-catenin downstream signaling pathways. CTGF has been found to mediate signaling during proliferative invasion, but its involvement in LUAD remains unclear (*28*).

**Figure 5.**
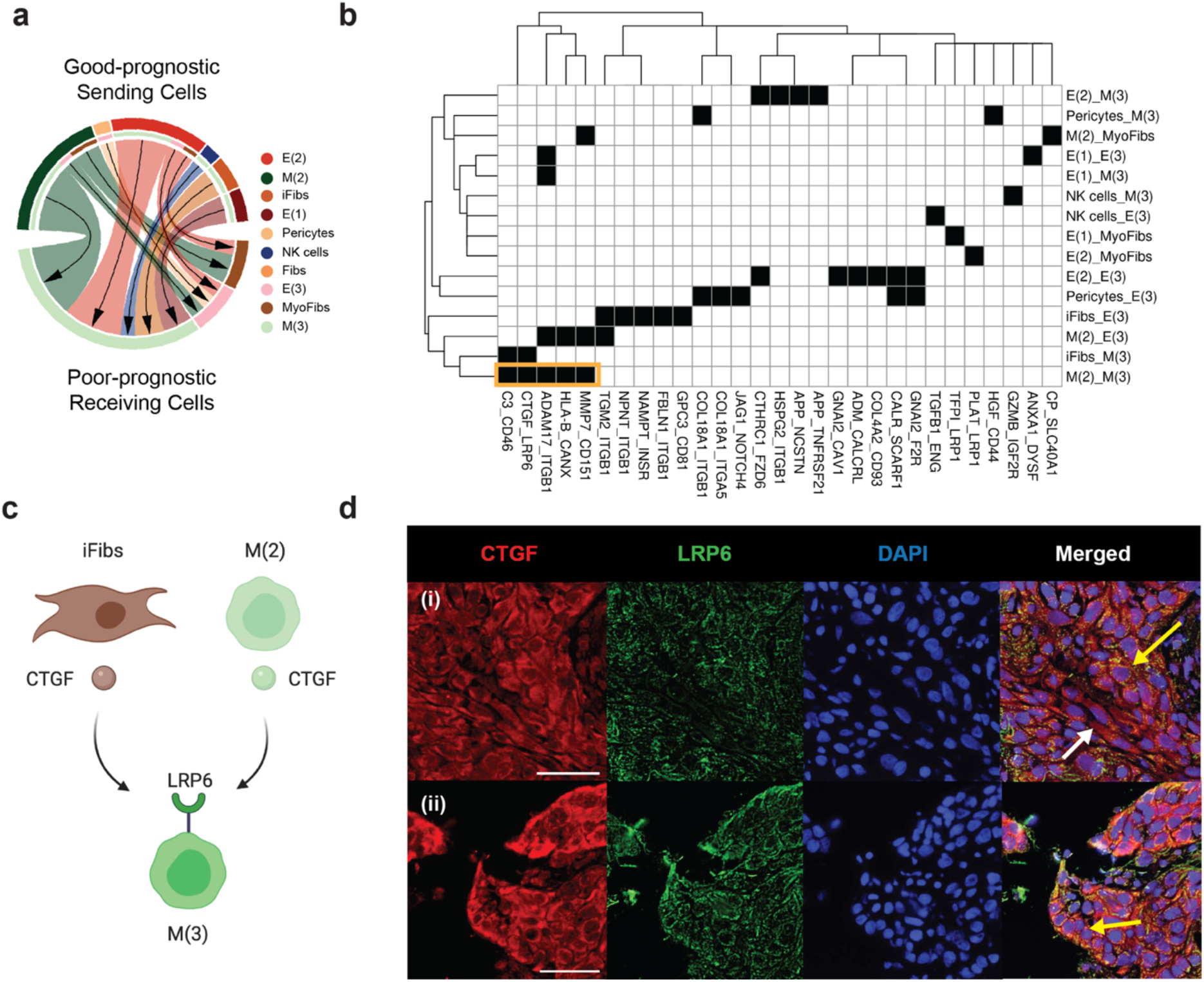
Paracrine-Autocrine CTGF:LRP6-axis detected in malignant subpopulations in high grade lung adenocarcinoma. **(a)** Circle plot indicates ratio of interactions occurring in mixed-prognostic subpopulations. The color of the chords corresponds to the legend on the right. **(b)** Binarized heatmap of cell-cell interactions between good-prognostic sending subpopulations and poor-prognostic receiving subpopulations. Interactions were filtered by average receptor expression > 0.5. Black indicates a cell-cell interaction exists between the genes and white indicates no cell-cell interaction exists between the genes. Yellow box highlights interactions occurring between M(2) and M(3) **(c)** Highlighting two interactions (iFibs_CTGF: M(3):LRP6 and M(2)_CTGF:M(3)_LRP6) within the scREMI community that CTGF and LRP6 from the malignant cell is in. Full community network is in Supplementary Figures. **(d)** Immunofluorescence (IF) staining analysis of CTGF and LRP6 in fresh frozen high grade lung adenocarcinoma tissues. Red indicates CTGF. Green indicates LRP6, Blue indicates DAPI. The white arrow points at fibroblast-like CTGF+LRP6-subpopulations. The yellow arrows in (i) and (ii) point at subpopulations with co-localization of CTGF and LRP6, indicating an interaction between the ligand-receptor pair. Yellow arrow in (ii) is pointing at a group of co-localized cells located at the edge of the tumor nest. Scale bars represent 50μm.

To determine the potential phenotypic role of CTGF:LRP6, we created a downstream signaling network for malignant cells using signaling pathway genes from KEGG and protein-protein interactions from the publicly available database BioGRID (*33, 34*). We set the edge-weight as the correlation between two signaling genes calculated using the bulk flow-sorted RNA-seq dataset and measured the log-transformed eigenvector centrality score (ranging from 0 – 1) of each node. A low-score indicates that the receptor has low correlation with its downstream pathway genes and a high-score indicates that the receptor has high correlation with its downstream pathway genes. This centrality score represents how influential a node is in the network in terms how correlated it is with each of its nodes are to the downstream network (Supp Fig 5e). *LRP6*’s downstream nodes were *GSK3B* and *CTNNB1* (*β*-catenin) with a centrality score of 0.27 and 0.31, respectively, which are relatively high given the mean centrality score of 0.05 (Supp Fig 5g – f). This relatively high centrality score indicates that the genes were correlated to their signaling neighbors within the downstream network. Within the scRNA-seq dataset, M(3) cells expressed *GSK3B* and *CTNNB1* and M(2) cells expressed *CTNNB1* (Supp Fig 5i). The expression patterns suggest that the paracrine interaction CTGF:LRP6 may activate WNT signaling downstream within M(3) to induce an invasive phenotype. Interestingly, previous studies identified a Lrp6-Gsk3b-*β*-catenin-Tcf-Ctgf autocrine axis within rat sarcomatoid mesothelioma, a type of lung cancer (*35*).

To confirm whether the co-localization of CTGF:LRP6 occurs in primary LUAD, we performed immunofluorescence (IF) imaging of CTGF, EpCAM, and PanCK in a high-grade lung adenocarcinoma tissue (Supp Fig 6a, i & ii). CTGF co-localized with the two epithelial markers (EpCAM, and PanCK), which suggests that CTGF is expressed by malignant subpopulations. CTGF was also expressed in regions of spindly cells that were negative for EpCAM and PanCK, which suggests expression in fibroblasts. Next, we performed IF imaging to simultaneously measure CTGF and LRP6 protein expression. In high-grade LUAD, we observed subpopulations of cells that exhibit either epithelial-like or fibroblast-like morphologies that may suggest they may be malignant or fibroblasts subpopulations, respectfully. CTGF co-localized with LRP6 within epithelial-like cells, whereas the fibroblast-like cells expressed solely CTGF (Fig 5d(i)). This is consistent with our inference that CTGF is expressed by both malignant and fibroblasts and interacts with LRP6 on malignant cells. We speculate that CTGF:LRP6 signaling transforms the M(2) cells into poorer-prognostic M(3) cells. Interestingly, we also observed that epithelial-like cells that were near the edge of the malignant nest had stronger co-localization signal for CTGF and LRP6 on the IF images (Fig 5d(ii)). We did not observe in CTGF:LRP6 co-localization low-grade LUAD.

## Discussion

REMI is a novel algorithm that utilizes graph-based approaches to generate a global network of cell-cell crosstalk by estimating conditionally-dependent LR pairs on high-dimensional data. As opposed to current methods, REMI captures the complexity of multi-cellular interactions and the potential confounding effects that LR pairs may have upon one another. Our approach offers the main advantage of identifying co-dependent interactions on a global scale with small sample size by leveraging prior knowledge, network analysis, and sparsity principles. We demonstrated REMI’s performance on a large transcriptomic dataset from TCGA, showing that estimating the inverse-correlation matrix using REMI is robust. We then sampled the TCGA data to create smaller datasets to quantify performance of REMI in terms of sensitivity and specificity. As the sample size decreased, we found that REMI’s specificity decreased less than its sensitivity, relative to other approaches, in terms of the “true” edges as derived from the full dataset. REMI’s higher specificity is favorable when choosing interactions for experimental verification in order to avoid false positives.

REMI is a modularized algorithm that allows investigators to adapt each component to best fit their biological question. In step 1, we used the FANTOM5 database as our reference for literature-supported LR pairs, but the database can be substituted or complemented with any other LR list. In step 2, we used the Louvain community detection algorithm, but this can be substituted for the investigator’s clustering algorithm of choice. REMI’s community detection step is hierarchical, which allows it to accommodate high-dimensional datasets while retaining high specificity. Thus, it can be applied to the many recent single-cell studies that have revealed substantial intra-tumoral heterogeneity. Finally, REMI was applied to transcriptomics data but it can be applied to proteomics data as well.

We applied REMI to a LUAD bulk flow-sorted RNA-sequencing dataset composed for four cell types (malignant, immune, fibroblast and endothelial) to re-assemble a highly-specific LUAD microenvironment interactome. We showed that REMI-LUAD captured tumor-stroma and tumor-stroma-immune specific communities that were enriched for ECM remodeling interactions and highlighted the prevalence of tumor-stroma-specific interactions. To expand the cell-type resolution of REMI-reconstructed networks, we projected a publicly available scRNA-seq data onto REMI-LUAD to construct scREMI-LUAD and uncovered potential interactions associated to progression of lung adenocarcinoma. The projection provided us with a higher resolution view of cell-type interactions. In particular, we observed that previously labeled autocrine signaling interactions within REMI-LUAD became paracrine signaling interactions between sub-populations of a given cell-type in scREMI-LUAD. This observation suggests that the currently documented autocrine signaling interactions may be involved in paracrine signaling between different cell states for a given cell type.

To infer a cancer progression signature within scREMI-LUAD, we calculated the prognosis of each cell sub-population accordingly to its signature patterns. We then applied unsupervised analysis of prognostic-specific cellular crosstalk signatures and ordered interactions based on prognosis as a proxy to tumor progression. Interactions involving good-prognostic subpopulations comprised of anti-tumoral or pre-invasive activities, whereas poor-prognostic subpopulations were involved with more invasive phenotypes. We reason that mixed-prognostic interactions may be associated with progression from a less to more invasive tumor microenvironment. Specifically, we experimentally validated a mixed-prognostic community of LR pairs (M(2)_CTGF:M(3)_LRP6 and iFibs_CTGF:M(3)_LRP6). Interestingly, anti-CTGF therapy is entering Phase 3 clinical trials for idiopathic pulmonary fibrosis (IPF) and shows promise in pancreatic cancer (*36*). This work suggests anti-CTGF therapy may be relevant to inhibit progression in LUAD and that other novel interactions found in scREMI-LUAD may also be potential therapeutic targets.

While REMI can identify highly-specific LR pairs, it has limitations. First, the utility of REMI may be limited by the assumption that interacting ligands and receptors are correlated. By leveraging these correlations between ligand and receptor genes representative of LR pairs, interactions that involve secretory molecules, such as chemokines which are typically secreted by immune cells, may not be captured well by REMI since chemokines can travel to distant tissue sites. Second, REMI does not yet account for large protein complexes. One way to potentially address this issue is to regularize edges between receptors in the same protein complex along with the LR pairs in the community. Third, REMI does not account for external influences on LR pairs, such as spatial or temporal effects. This may be remedied by implementing a weighted graphical Gaussian model instead. A weighted approach may also allow users to capture LR affinity or downstream signaling effects. Fourth the significance test is designed for each community since the edge permutations changes the communities identified by REMI. This stochasticity is difficult to capture in REMI’s current statistical test. Calculating p-values for all edges within the network is also time-intensive. Further work needs to be done to implement a fast interactome-wide test for statistical significance.

In summary, we developed REMI and demonstrated the benefit of using REMI networks as a discovery tool. We took advantage of the robustness of bulk RNA-seq data that captures the power obtained from analyzing the homogeneity within tumor, stroma, and immune cell subpopulations, and used the heterogeneous properties of scRNA-seq to further elucidate the microenvironment interactome. Our results provide a novel way to infer global cellular crosstalk for subsequent functional validation. We anticipate that REMI can be applied to many more tissue microenvironments to better understand normal and diseased processes and develop more precise therapeutics.

## Materials and Methods

### Bulk flow-sorted RNA-seq data

We used bulk flow-sorted RNA-Seq data (GSE111907) on 40 tumor samples that were sorted using flow cytometry into the four cell types of interest: immune, endothelial, fibroblast, and malignant cancerous cells using the markers CD45+/EpCAM-, CD31+/CD45-/EpCAM-, CD10+/EpCAM-/CD45-/CD31-, and EpCAM/CD45-respectively. Patients without data from all four cell-types were removed. The data included bulk RNA-sequencing measurements for each cell type as defined as *X* = {*X_c_*_1_, *X_c_*_2_, *X_c_*_3,_ *X_cn_*}. The TPM values were log-transformed (log2(TPM+1)) and standardized to mean 0 and standard deviation 1.

### Publicly available databases

*FANTOM5 database.* We obtained a total of 1,904 validated ligand-receptor pairs (650 ligands and 594 receptors) from the FANTOM5 database (Ramilowski et al. 2015). The database extracted known ligand-receptor pairs from the Database of Ligand-Receptor Partners (DLRP), IUPHAR, and the Human Plasma Membrane Receptome (HPMR). We filtered for LR pairs that had literature-supported evidence and were experimentally validated. We appended 12 more literature-supported experimentally-validated LR pairs not found in the database: PDL1/PD-1, PD-2/PDL1, CD80/CTLA4, CD80/CD28, CD86/CD28, GREM1/KDR, PDL2/PD-1, NECTIN2/CD226, NECTIN2/TIGHT, PVR/TIGHT, SIGLEC1/SPN, CTGF/TNFRSF1A. *KEGG Database.* From the Kyoto Encyclopedia of Genes and Genomes (KEGG) database, we downloaded all the pathway genes from the Environmental Information Processing category. This includes signaling pathways associated with membrane transport, signal transduction, and signaling molecules and interactions. Genes from the phosphotransferase system (PTS) and bacterial secretion system were excluded. *BioGRID Database.* We downloaded the *Homo Sapiens* proteins from BioGRID and their interaction network. Only protein-protein interactions that were experimentally validated were used. We removed proteins that were found in the publicly available database CRAPome (*37*). The remaining BioGRID network has 18074 proteins and 176120 interactions.

### LTMI reconstruction

For the LTMI reconstruction, the log-transformed bulk flow-sorted RNA-seq data from Gentles et al. was split into lung adenocarcinoma and lung squamous cell carcinoma patients. Using only the lung adenocarcinoma patients (n=17), ligand and receptor genes were filtered by TPM > 10. A graph was constructed by creating a node for every ligand and receptor gene that passed the filtering criteria and an edge was drawn if it was a known pair according to FANTOM5 database. This was cross-referenced with the LTMI built on all samples. The igraph R package was used to construct and illustrate the graph. We also used igraph to identify components in the network. To calculate partial correlation for the subgraph, we used the following equation:

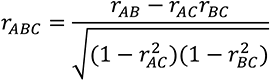

This equation was extended to capture all combinations of gene interactions within the subgraph. Correlation was measured using the gene expression dataset.

### REMI algorithm

***(i) Build weighted bipartite LR network.*** Using a user-inputted LR list, a LR network is created. The default LR list is obtained from the FANTOM5 database. Cell-types from the input dataset are denoted as *c*_1_, c_2_, …,*c_n_*∈*X*. *x_c_* represents all ligands found in cell-type *c*(*x*_1_,*x*_2_,…,*x_a_*∈*X_c_*). *y_c_* represents all receptors found in cell-type *c*(*y*_1_,*y*_2_,…,*y_b_*∈*X_c_*}). Given dataset *X*, we represent all potential LR pairs interacting between all cell-types in the undirected network *G_LR_* = (*V*_1_=*V_x_*∩*V_y_*,*E*_1_) where *V_x_* = {*x*:*x_c_1__*,*x_c_2__*,…,*x*_c_n__} and *V_y_* = {*y*:*y_c_1__*,*y_c_2__*,…,*y*_c_n__}. An edge represents a relationship between ligand and its cognate receptor denoted as *E*_1_ ={(*x*,*y*):*x*∈*V_x_*,*y*∈*Vy, (x,y)*∈*N_FANTOM5_*}. Edge weight *w_ij_* in *G_LR_* is the Pearson correlation between *x_i_* and *y_j_* calculated using the gene expression data. ***(ii) Detect cognate LR communities.*** Communities (*C_k_*) are identified in *G_LR_* using Louvain community detection algorithm. The algorithm identifies clusters of nodes, called communities, that are optimized for maximum modularity (*Q*) within the network:

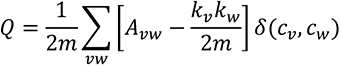

*A_vw_* is the Spearman correlation between nodes *v* and *w*. *m* is the number of edges. *k* is the degree of node. δ is the Kroenecker delta piecewise function indicating whether node *v* and *w* are in the same community. *c* is an indicator function for whether a node is in a community. The measurement ensures the communities detected are data-driven and not based on chance. *G_LR_* can contain disjoint components since some nodes were removed due to their low-expression within the dataset. Disconnected small components are referred to as a community. If the number of nodes within *C_k_* is greater than the sample size, we perform hierarchical community detection using Louvain to further reduce the density of the communities until the number of nodes in every communities is less than or equal to the sample size. We then iterate between every pairwise combination of communities and create a pseudo-community, or between-community. The between-communities contains nodes that have an edge between the two communities.

### (iii) Remove conditionally independent LR pairs

According to the Hammersley-Clifford theorem, an inverse covariance matrix, or partial correlation, can be used to indicate relationships between nodes. A zero in an inverse covariance matrix indicates conditional independence between the two variables in the model given all others. The algorithm graphical lasso (GLasso) utilizes this concept and estimates the inverse covariance matrix that best fits the observations in a multivariate Gaussian distributed dataset (*17*). A *l*1 penalty is added to the log-likelihood of a Gaussian Markov Field to add data-driven sparsity to the model. Here is the optimization function:

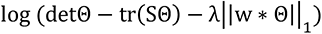

In this equation, Θ = *σ*^−1^ and S is the empirical covariance matrix. *w* represents the weighted adjacency matrix. *tr* is the trace of the matrix or the sum of the diagonal. ||Θ||_1_ is the *l*1 penalization of the inverse covariance matrix. When λ increases, the network becomes sparser. When λ is zero, the resulting network is equivalent to a partial correlation network (*17*). We utilize GLasso from the glasso R package to measure the conditional dependence between two connected nodes while controlling for the effect of all other nodes for each community detected. Since the communities identified within the largest network component are not disjoint clusters, we also created graphical models using the nodes and edges between communities. These between communities only include nodes that have edges between communities and excludes nodes that do not. Communities with one node were not included.

The GLasso similarity matrix can be represented as a fully connected LR community graph. This includes all potential edges between ligand and receptors (LR), ligand and ligand (LL), and receptor and receptor (RR). We simplify the LR prediction problem by focusing on only LR interactions and make the assumption that the underlying network that represents the data only contains edges between ligands and receptors. LL and RR edges within the *G_LR_* communities are penalized and set to zero in the weighted adjacency matrix, indicating conditional independence. The penalized weighted matrix is defined as:

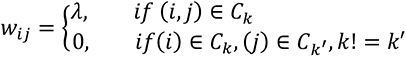

The λ tuning parameter in the penalized log-likelihood equation is chosen by iterating through 20 values logarithmically spaced between lambda values 0.01 to 0.9. The optimal *λ* is the one that best minimizes the Bayesian information criterion (BIC). The accuracy of the graphical lasso method is dependent on the number of features in the network. ***(iv) Reconstruct communities into predicted interactome*.** The final reconstructed LR network includes edges between nodes that have a weight larger than zero. ***(vi) Significance test for p-*value.** We provide an optional additional tool to calculate p-values for an edge in a community. P-values are calculated by measuring the occurrence of a test statistic compared to its null. In the REMI algorithm, a LR pair is a subset of a test statistic, the regularized covariance matrix. Therefore, our significance test measures the conditional probability that a LR pair is conditionally dependent within its REMI community given added randomization. First, we sample from a uniform distribution of correlation values and replace the correlation value of the LR pair of interest. Then, REMI is re-run 1000 times to generate a distribution of predicted scores for the LR pair. A regression tree is fit on the sampled correlation values to predict binary REMI scores. A null distribution is then generated by simulating a null training dataset by generating a Gaussian distribution (*<u>G</u>*) and training it on the regression tree to predict a null set of outcomes (*w*). *w* is multiplied by the Wilshart distribution to create a null distribution that represents the conditional test statistic (*38*). The p-value is then calculated by using the LR pair’s data-derived correlation value (*R*[*i*,*j*])). The first equation represents the one-sided p-value and the second equation represents the two-sided p-value.

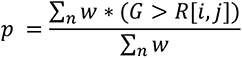

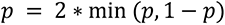

### Validation dataset

We tested the method on a bulk RNA-Sequencing dataset downloaded from the publicly available database, the Cancer Genome Atlas (TCGA). The NSCLC RNA-Sequencing dataset contains bulk RNA-Sequencing data for 1013 NSCLC patients. This includes 584 adenocarcinoma and 429 lung squamous cell carcinoma patients. The data was log-transformed and scaled to mean of 0 and variance of 1.

### Simulations

The “gold standard” interactome was generated by running GLasso on *G_LR_* generated using LR genes found in the TCGA NSCLC bulk RNA-sequencing dataset (*V* = 893, *E* = 1331). A true positive edge is an edge with weight greater than zero. All other edges are true negative edges. Five cohort sizes (*n* = 10, 25, 50, 100, and 2000) were sampled 50 times each from the whole TCGA NSCLC cohort to simulate various cohort sizes. REMI was run on each sampled cohort dataset using default parameters and unfiltered list of ligand and receptor genes. GLasso was also performed on each sampled cohort dataset with restrictions on the LL and RR edges. Performance metrics were calculated by setting the gold standard interactome as the true network. The same analysis was performed by separating within and between communities’ predictions and calculating their performance metrics. Noise was also added to the sampled datasets and the performance metrics were calculated.

### Comparison against other methods

*NicheNet.* We ran NicheNET on the TCGA NSCLC bulk RNA-sequencing dataset using the same filtered genelist as the ones used in the TCGA simulations described above. The background geneset used were the genes found in the ligand target matrix of the receiving cell. The LR list was replaced with the FANTOM5 list we generated. For TCGA data, we treated the bulk dataset as an autocrine signaling. We ran NicheNET on every autocrine and paracrine combination of cell types. We used the best ligand filter of Pearson > 0. *NATMI.* We used both log-transformed and scaled log-transformed TCGA RNA-sequencing dataset as input and set all samples as one cell type. Default parameters were used and edges were filtered based on specificity score. *CellphoneDB*. We used the log-transformed TCGA RNA-sequencing dataset as input for the method. We used the database provided and filtered for significant LR pairs based on the significant means score. *CCCExplorer.* We ran CCCExplorer on the TCGA NSCLC bulk RNA-sequencing dataset and bulk flow-sorted RNA-seq dataset. To identify DEG ligands for the input, we used samr to identify differentially expressed genes (DEG). For TCGA, we identified DEG between healthy and lung cancer patients. For the bulk flow-sorted cohort, we identified DEG between the signaling cell and receiving cell. The DEG cut-offs used were logFC > 1.2 and q-val < 0.05. The same DEG input from NicheNET was used. We calculated cell-cell interactions for every pairwise paracrine and autocrine combination.

### REMI on LUAD

For the REMI interactome construction, the log-normalized gene expression measurements were binned into three evenly split discretized groups for each cell type. Genes with expression levels falling within the first group, representing low expressed genes, were removed. The dispersion ratio (variance/mean) was calculated for each gene and normalized within each bin. Genes in the first bin, which includes genes with negative and fairly low expression levels, were removed. The data was scaled after filtering and the filtered dataset was used to identify ligand and receptor genes. The scaled unfiltered gene expression dataset was used for all other analysis. All other parameters in REMI were set to default.

### Gene enrichment analysis

The clusterProfiler R package was used to perform gene enrichment analysis. The genelists were filtered for the databases: Hallmark, BIOCARTA, and REACTOME. Enrichments with adjusted p-value < 0.05 were used for analysis.

### scRNA-seq analysis

*Data* We used publicly available single-cell RNA-Seq datasets E-MTAB-6149 and E-MTAB-6653 generated by Lambrecht et al (*21*). The droplet-based scRNA-Seq (10X genomics) datasets comprised of two squamous cell carcinoma patients, two adenocarcinoma patients, and one non-small cell lung carcinoma patient. We used the Seurat V3 R package to process the lung adenocarcinoma carcinoma patients and used pre-defined cell type labels defined by the authors (*39*). *Preprocessing*. Cells with more than 10% mitochondria content, over 6000 and under 100 features were filtered out. We used the top 30 principal components and the IntegrateData function in Seurat to correct for batch effect between the patients. Malignant, fibroblast, and endothelial cells were re-clustered using Louvain clustering on a Shared Nearest Neighbor (SNN) graph with 10 principal components using 0.2, 0.2, and 0.1 resolution respectively. Differentially expressed genes across all cell types and within clusters were calculated using MAST with default parameters and normalized counts (averaged log2FC > 0.25, Bonferroni corrected p-val < 0.05) (*40*). Visualizations are shown using UMAP (30 principal components). PHATE was used to infer the underlying hierarchical manifold in each cell-type using default parameters (*41*). Each cell-type population was imputed using ALRA after identifying subpopulations (*42*).

### Deconvolving interactome

LR genes in the REMI LUAD interactome were filtered based on whether they were expressed in their respective cell-type (average expression > 0.4). The cell-type metadata associated with each node in the interactome was relabeled using the subpopulation the ligand or receptor was found in. The cell-type must be the same between bulk and single-cell data. Nodes with genes that were found across multiple subpopulations within the same cell type were duplicated and an edge is drawn to its respective predicted cognate pair.

### Measuring prognosis of subpopulations

Bulk LUAD-specific prognosis scores were downloaded from the PRECOG metaZ database. Then, the normalized log-transformed average expression level for the ligands within each sub-population were calculated using the ALRA imputed gene expression data. Each subpopulation was imputed independently. For each subpopulation, ligands that were not differentially expressed (DE), calculated by MAST in prior step, was removed. DE ligand cognate receptors with an average scaled expression less than 0 were removed. Subpopulations with less than three DE ligands predicted to be secreted were also removed from the calculations. In the remaining list of LR pairs, we then multiplied the differentially expressed ligand’s averaged expression level by its PRECOG score (*a* = 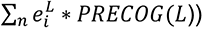. The total gene expression values of ligands 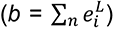 were aggregated and the prognosis of the subpopulation by *a*/*b* was calculated. Additional filtering of the LR pairs was then applied on duplicate LR pairs that have the same genes, but different sending and receiving cells. The pair with the closest PRECOG score between sending and receiving cells was kept and the rest were filtered out.

### Immunofluorescence and H&E staining

Fresh frozen human lung adenocarcinoma tissues were serially sectioned at a thickness of 8 μm and mounted on poly-L-lysine-coated coverslips. One section was successively fixed in 95% ethanol and 10% formalin for 10 minutes each and subjected to progressive H&E staining following the standard protocol described (*43*) for pathological review by a pathologist. For IF analysis, sections were fixed in acetone for 10 minutes at room temperature (RT) and then washed twice with PBS. Fluorophore bleaching was performed at RT for 90 minutes to reduce autofluorescence by immersing the fixed coverslip in freshly prepared bleaching solution (4.5% (wt/vol) H_2_O_2_ and 20 mM NaOH in PBS) and illuminating the container between two LED light panels, followed by four washes with PBS to remove bleaching solution (*44*). The pre-treated coverslips were blocked with 1% horse serum in PBS for 30 minutes, and incubated with primary antibodies in incubation solution (1% bovine serum albumin, 1% normal donkey serum, 0.3% Triton X-100, and 0.01% sodium azide in PBS) in a humidified chamber overnight at 4°C. After three washes in PBS for 15 minutes per wash, the coverslips were incubated with secondary antibodies in incubation solution for 1 hour at RT in dark. The coverslips were intensively washed with PBS and then stained with DRAQ5 (Thermo) at 1: 1000 in PBS for 10 minutes. The sections were observed under the BZ-X800 fluorescence microscope (Keyence, IL, USA). Antibodies used in the study include anti-CTGF (ab6992, Abcam), anti-LRP6 (MAB1505, R&D System), anti-EpCAM (#2929, Cell Signaling Technology), anti-pan Cytokeratin (ab86734, Invitrogen), Cy3-conjugated goat anti-rabbit IgG (ab6939, Abcam), goat anti-mIgG1 Alexa Fluor 488 (A-21121, Invitrogen) and donkey anti-mouse IgG H&L Alexa Fluor 488 (ab150105, Abcam) antibodies.

### Downstream influence score

An undirected downstream signaling pathway network *G_ds_*= (*V*_2_,*E*_2_) is created for each cell type in the dataset. Each *G_ds_* consists of approximately 1000 nodes representing signal transduction pathway genes (*N_KEGG_*) downloaded from the publicly available database, Kyoto Encyclopedia of Genes and Genomes (KEGG) (*V*_2_ = {*x*:*x*∈*N*_KEGG_}) (*34*). An edge between two nodes represents a physical protein-protein interaction between the nodes, as indicated in the database, BioGRID (*E_2_* = {(*x_i_*,*x*_j_)∈*N_BioGRID_*}) (*33*). The edge weight in the network is set as the Pearson correlation between the two gene nodes calculated from the gene expression data from the cell type of interest. For each *G_2_* per cell type, we calculate the eigenvector centrality for each node. Eigenvector centrality is a centrality measurement of a node *x_i_* that is relational to the sum of importance of its neighbor nodes: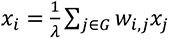, where *w_i,j_* is the weight between nodes *x_i_* and *x_j_*. The centrality measurement for each receptor within *G_ds_* is the receptor’s influence score.

## Acknowledgements

We gratefully acknowledge L.K. for the discussions on cell states in lung cancer, G.B. for discussions on experimental validation, C.G. for helping us resolve issues with autofluorescence in the IF images, and B.A., D.K., W.Z., and A.B. for the discussions on the methodology. We also thank A.G., P.M., and T.D. for discussions about the biological properties of ligand-receptor signaling and method development. A.Y is supported by the National Library of Medicine grant TL15LM007033, Tobacco-Related Disease Research Program pre-doctoral fellowship grant T29DT0760, and NIH U544CA209971. S.K.P is supported by NIH U54CA209971.

## Author Contributions

A.Y. and S.K.P. designed the methods and spearheaded the direction of the methods and manuscript. Y.L. conducted all experimental validation. Y.L. and I.L. contributed to experimental validation design. C.Y. and A.E.C. contributed to computational analysis. M.G.O. reviewed H&E slides. J.T. designed and contributed to method’s statistical significance test.

## Data Materials and Availability

All will be available upon request and after publication.

## References

1. S. K. Wculek, I. Malanchi, Neutrophils support lung colonization of metastasis-initiating breast cancer cells. Nature. 528, 413–417 (2015).

2. E. Armingol, A. Officer, O. Harismendy, N. E. Lewis, Deciphering cell–cell interactions and communication from gene expression. Nat. Rev. Genet. (2020), doi:10.1038/s41576-020-00292-x.

3. M. Efremova, M. Vento-Tormo, S. A. Teichmann, R. Vento-Tormo, CellPhoneDB: inferring cell-cell communication from combined expression of multi-subunit ligand-receptor complexes. Nat. Protoc. 15, 1484–1506 (2020).

4. R. Hou, E. Denisenko, H. T. Ong, J. A. Ramilowski, A. R. R. Forrest, Predicting cell-to-cell communication networks using NATMI. Nat. Commun. 11, 5011 (2020).

5. H. Choi et al., Transcriptome analysis of individual stromal cell populations identifies stroma-tumor crosstalk in mouse lung cancer model. Cell Rep. 10, 1187–1201 (2015).

6. R. Browaeys, W. Saelens, Y. Saeys, NicheNet: modeling intercellular communication by linking ligands to target genes. Nat. Methods. 17, 159–162 (2020).

7. A. Patsialou et al., Invasion of human breast cancer cells in vivo requires both paracrine and autocrine loops involving the colony-stimulating factor-1 receptor. Cancer Res. 69, 9498–9506 (2009).

8. L. Hernandez et al., The EGF/CSF-1 paracrine invasion loop can be triggered by heregulin beta1 and CXCL12. Cancer Res. 69, 3221–3227 (2009).

9. M. P. Kumar et al., Analysis of Single-Cell RNA-Seq Identifies Cell-Cell Communication Associated with Tumor Characteristics. Cell Rep. 25, 1458–1468.e4 (2018).

10. A. J. Gentles et al., A human lung tumor microenvironment interactome identifies clinically relevant cell-type cross-talk. Genome Biol. 21, 107 (2020).

11. J. E. Allen, W. S. El-Deiry, Regulation of the human TRAIL gene. Cancer Biol. Ther. 13, 1143–1151 (2012).

12. M. de Looff, S. de Jong, F. A. E. Kruyt, Multiple interactions between cancer cells and the tumor microenvironment modulate TRAIL signaling: implications for TRAIL receptor targeted therapy. Front. Immunol. 10, 1530 (2019).

13. D. Deng, K. Shah, TRAIL of hope meeting resistance in cancer. Trends Cancer (2020), doi:10.1016/j.trecan.2020.06.006.

14. T. Hartwig et al., The TRAIL-Induced Cancer Secretome Promotes a Tumor-Supportive Immune Microenvironment via CCR2. Mol. Cell. 65, 730–742.e5 (2017).

15. J. A. Ramilowski et al., A draft network of ligand-receptor-mediated multicellular signalling in human. Nat. Commun. 6, 7866 (2015).

16. K. Komurov, Modeling community-wide molecular networks of multicellular systems. Bioinformatics. 28, 694–700 (2012).

17. J. Friedman, T. Hastie, R. Tibshirani, Sparse inverse covariance estimation with the graphical lasso. Biostatistics. 9, 432–441 (2008).

18. X. Guo et al., Global characterization of T cells in non-small-cell lung cancer by single-cell sequencing. Nat. Med. 24, 978–985 (2018).

19. J. Winkler, A. Abisoye-Ogunniyan, K. J. Metcalf, Z. Werb, Concepts of extracellular matrix remodelling in tumour progression and metastasis. Nat. Commun. 11, 5120 (2020).

20. V. Gkretsi, T. Stylianopoulos, Cell adhesion and matrix stiffness: coordinating cancer cell invasion and metastasis. Front. Oncol. 8, 145 (2018).

21. D. Lambrechts et al., Phenotype molding of stromal cells in the lung tumor microenvironment. Nat. Med. 24, 1277–1289 (2018).

22. R. Kalluri, The biology and function of fibroblasts in cancer. Nat. Rev. Cancer. 16, 582– 598 (2016).

23. A. J. Gentles et al., The prognostic landscape of genes and infiltrating immune cells across human cancers. Nat. Med. 21, 938–945 (2015).

24. R. Kong et al., Myo9b is a key player in SLIT/ROBO-mediated lung tumor suppression. J. Clin. Invest. 125, 4407–4420 (2015).

25. R. K. Gara et al., Slit/Robo pathway: a promising therapeutic target for cancer. Drug Discov. Today. 20, 156–164 (2015).

26. R. Sadej, A. Grudowska, L. Turczyk, R. Kordek, H. M. Romanska, CD151 in cancer progression and metastasis: a complex scenario. Lab. Invest. 94, 41–51 (2014).

27. N. K. Altorki et al., The lung microenvironment: an important regulator of tumour growth and metastasis. Nat. Rev. Cancer. 19, 9–31 (2019).

28. A. L. Parker, T. R. Cox, The role of the ECM in lung cancer dormancy and outgrowth. Front. Oncol. 10, 1766 (2020).

29. Z. Chen et al., Connective tissue growth factor: from molecular understandings to drug discovery. Front. Cell Dev. Biol. 8, 593269 (2020).

30. A. Resovi et al., CCN-Based Therapeutic Peptides Modify Pancreatic Ductal Adenocarcinoma Microenvironment and Decrease Tumor Growth in Combination with Chemotherapy. Cells. 9 (2020), doi:10.3390/cells9040952.

31. J. Raisch, A. Côté-Biron, N. Rivard, A Role for the WNT Co-Receptor LRP6 in Pathogenesis and Therapy of Epithelial Cancers. Cancers (Basel). 11 (2019), doi:10.3390/cancers11081162.

32. K. E. Lipson, C. Wong, Y. Teng, S. Spong, CTGF is a central mediator of tissue remodeling and fibrosis and its inhibition can reverse the process of fibrosis. Fibrogenesis Tissue Repair. 5, S24 (2012).

33. C. Stark et al., BioGRID: a general repository for interaction datasets. Nucleic Acids Res. 34, D535–9 (2006).

34. M. Kanehisa, Y. Sato, M. Furumichi, K. Morishima, M. Tanabe, New approach for understanding genome variations in KEGG. Nucleic Acids Res. 47, D590–D595 (2019).

35. L. Jiang et al., Connective tissue growth factor and β-catenin constitute an autocrine loop for activation in rat sarcomatoid mesothelioma. J. Pathol. 233, 402–414 (2014).

36. G. Sgalla, C. Franciosa, J. Simonetti, L. Richeldi, Pamrevlumab for the treatment of idiopathic pulmonary fibrosis. Expert Opin. Investig. Drugs (2020), doi:10.1080/13543784.2020.1773790.

37. D. Mellacheruvu et al., The CRAPome: a contaminant repository for affinity purification-mass spectrometry data. Nat. Methods. 10, 730–736 (2013).

38. J. Markovic, J. Taylor, J. Taylor, Inference after black box selection.

39. T. Stuart et al., Comprehensive Integration of Single-Cell Data. Cell. 177, 1888–1902.e21 (2019).

40. G. Finak et al., MAST: a flexible statistical framework for assessing transcriptional changes and characterizing heterogeneity in single-cell RNA sequencing data. Genome Biol. 16, 278 (2015).

41. K. R. Moon et al., Visualizing structure and transitions in high-dimensional biological data. Nat. Biotechnol. 37, 1482–1492 (2019).

42. G. C. Linderman, J. Zhao, Y. Kluger, Zero-preserving imputation of scRNA-seq data using low-rank approximation. BioRxiv (2018), doi:10.1101/397588.

43. R. D. Cardiff, C. H. Miller, R. J. Munn, Manual hematoxylin and eosin staining of mouse tissue sections. Cold Spring Harb. Protoc. 2014, 655–658 (2014).

44. Z. Du et al., Qualifying antibodies for image-based immune profiling and multiplexed tissue imaging. Nat. Protoc. 14, 2900–2930 (2019).

